# Data Driven Disease Dynamics Models

**DOI:** 10.1101/2025.08.01.668106

**Authors:** Polina Banushkina, Sergei Krivov

## Abstract

Models that explicitly consider the dynamic nature of disease progression promise a more comprehensive analysis of longitudinal datasets and disease characterization. This paper presents a new framework that uses optimal reaction coordinates (RCs) to describe disease progression as a diffusion on a free energy landscape. This method addresses key challenges, including the curse of dimensionality, irregular sampling, and data imbalance, providing a theoretically optimal representation of stochastic disease dynamics. Additionally, we introduce a new validation criterion that outperforms traditional metrics like AUC in distinguishing between optimal and sub-optimal RCs. The proposed framework provides a feasible approach for constructing datadriven models of disease dynamics from irregular, real-world longitudinal clinical datasets.

## 1 Introduction

Artificial intelligence (AI) research in the analysis of longitudinal data offers significant benefits to the biomedical field, particularly in healthcare. A review of recent biomedical and clinical applications shows promising results across a wide range of medical specialties [1, 2, 3]. With the right tools, analyzing clinical patient data can help predict diseases early, enabling timely intervention for high-risk individuals. Despite numerous studies demonstrating AI’s potential in healthcare, successful implementations in medical practice remain limited [4]. Several works explore the main challenges and limitations of AI in the biomedical field [1, 2, 3, 4] and emphasize the need for quality criteria in AI-based prediction models for healthcare applications [4, 5].

Models that consider disease dynamics could offer distinct advantages in analyzing longitudinal clinical datasets, such as electronic health records (EHR). For example, they demonstrate increased robustness when handling censored data - cases where individuals drop out of studies or do not experience the event of interest during the study period - a common challenge that complicates standard modeling approaches [6, 7, 8]. Additionally, these models simplify the definition of surrogate end states by eliminating the need to define them individually for each patient. While machine learning (ML) algorithms show considerable promise for longitudinal data analysis, current approaches predominantly rely on non-dynamic, static supervised classification models [1]. In their comprehensive review, Cascarano et al. highlighted this limitation, noting a significant “lack of algorithms exploiting the dynamic aspect of longitudinal datasets” [1].

Ideally, a model should capture the complete dynamics of disease progression, from healthy to diseased states, with particular emphasis on the critical transition phase. This period, when individuals are at elevated risk but still have normal laboratory results, presents the optimal window for early intervention. Early detection at this stage is crucial for implementing preventive measures before the disease manifests. We argue that the full potential of longitudinal datasets can be realized in data-driven models of disease dynamics.

To clarify what we mean by such models - since the concept of a dynamical disease model can vary - we describe what we consider an “ideal” model of disease dynamics. The most comprehensive description of such dynamics can be achieved using finite-state Markov chains [9, 10]. In this framework, disease progression is modeled as a Markov (memory-less) stochastic process, where an organism’s future behavior is determined probabilistically by its current state - defined by the complex of genome, proteome, metabolome, epigenome, age, environment, and all other relevant factors (hereafter the “configuration space”). Using an ensemble of longitudinal patients EHR trajectories, one can construct the transition probability matrix, which describes the probabilities of transitions between different micro-states within this configuration space. This matrix provides a complete description of patient and disease dynamics, enabling various prognostic applications. For instance, given a patient’s current conditions, one can generate trajectories that accurately predict future developments. By sampling and averaging these trajectories, one can compute key clinical characteristics, such as: likelihood of disease onset, mean time to disease development, probability of disease onset within a specific time intervals (e.g., 24 hours).

However, the practical implementation of such a brute-force model of disease dynamics, faces several fundamental challenges, some of which are shared with application of other ML techniques in the analysis of EHRs:

### Curse of dimensionality

The statistics required to construct the Markov chain grow exponentially with the dimensionality of the configuration space. This increase in dimensionality arises from the need to include numerous variables to fully characterize a patient’s state and keep the dynamics Markovian.

### Highly irregular sampling

EHRs typically contain missing values and exhibit irregular time intervals between measurements, posing challenges for conventional ML methods that require complete, regularly spaced data. While preprocessing techniques such as data imputation and polynomial interpolation have been proposed [10, 1], evidence suggests that some approaches may fail to effectively capture missing patterns, leading to sub-optimal analysis [11, 1].

### Imbalanced sampling

The statistical rarity of disease events within populations creates significant data imbalance. This imbalance reduces their contribution to training loss functions, resulting in models that excel at describing healthy dynamics but perform poorly in capturing disease transitions.

### Evaluation metrics

Clinical implementation demands rigorous model evaluation [5]. While the Area Under the Curve (AUC) of the Receiver Operating Characteristic (ROC) is widely used, its performance deteriorates with imbalanced datasets [1, 12, 13]. This limitation underscores the necessity of alternative or complementary evaluation metrics specifically tailored to healthcare applications [14, 4].

### Visualization and explainability

The “black box” nature of AI models presents significant challenges in visualizing and interpreting the learned disease dynamics encoded in the transition probability matrix. Enhancing interpretability is essential for clinical adoption.

Given the limitations of existing approaches and the challenges outlined above, there remains a critical need for rigorous, data-driven dynamical approaches. These approaches should effectively analyze longitudinal EHR data to build models of disease dynamics while being practical for medical implementation. To address these challenges, we propose a practically feasible approach for constructing models of disease dynamics from longitudinal datasets. This approach is based on the framework of optimal reaction coordinates, preserving the advantages of dynamical modeling while avoiding the limitations of the brute-force method outlined above.

Briefly, instead of using discrete Markov chains, we describe disease progression as diffusion on a free energy landscape parameterized by optimal reaction coordinates (RCs). The optimal RCs are determined from input variables or features (e.g., longitudinal biomarker panels). The coordinate(s) measure progress over time, providing the best possible description of the complex stochastic process capturing the entire disease dynamics of transitioning from a healthy to a diseased state. Originally, this approach was specifically designed for accurately describing the dynamics of rare events, such as protein folding. It has been successfully applied to longitudinal atomistic protein folding trajectories [15, 16, 17, 18], as well as to other domains [19, 9]. Here, we extend this approach to non-equilibrium, imbalanced data with irregular sampling. By optimally projecting complex dynamics data onto one or a few RCs, this approach circumvents the curse of dimensionality while preserving key dynamical properties of interest. It is particularly well-suited for processing highly irregular longitudinal datasets typical of clinical data, with an emphasis on capturing rare disease dynamics rather than predominantly healthy patient states. Its stringent validation criterion ensures robust performance on these challenging datasets. Moreover, the overall disease dynamics is intuitively represented as diffusion on a free energy landscape, enhancing interpretability and enabling a data-driven definition of disease states as metastable regions or free energy basins. Finally, the non-parametric construction of optimal RCs eliminates the need to select an appropriate neural network architecture, simplifying deployment. In summary, this new framework represents a robust, versatile, and clinically relevant tool for analyzing disease dynamics, opening new possibilities for early diagnosis and medical intervention. Fig. 1 presents the workflow illustrating how the optimal RC framework is applied to analyze disease dynamics.

**Figure 1:**
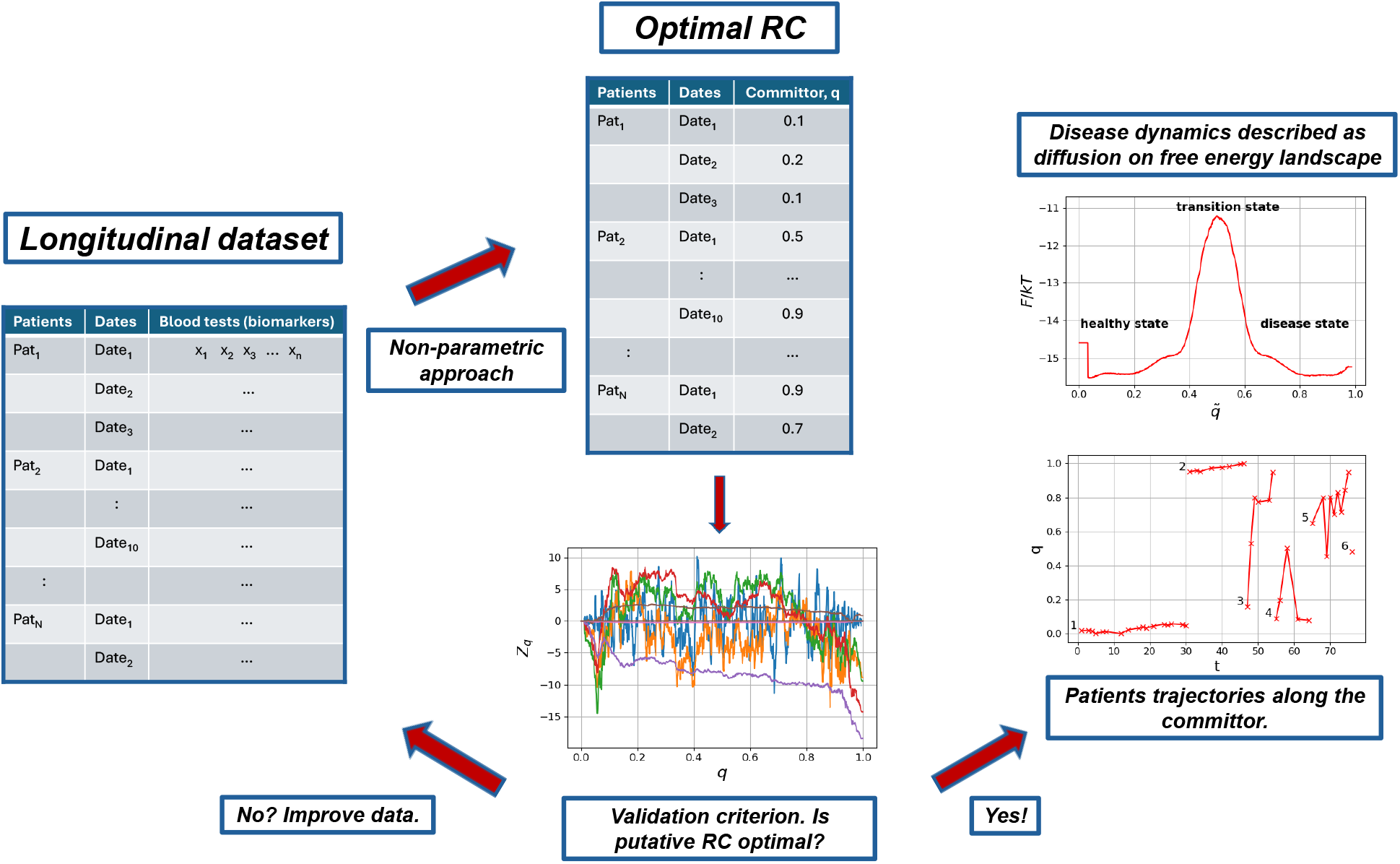
Workflow diagram illustrating the application of the proposed framework to clinical data analysis. Patient records (EHR) are combined into a single longitudinal dataset. The non-parametric approach is applied to estimate the committor time series, i.e., the committor value for each observation for every patient. The validation criterion is then applied to check whether the committor estimate is sufficiently close to the true value. If not, the dataset can be expanded with additional biomarkers. If yes, the obtained committor time series can be used to describe disease/patient dynamics. For example, it can be used to construct a free energy landscape that represents overall disease dynamics as diffusion on a free energy surface, where free energy basins provide a data-driven definition of disease states and the transition state marks the critical phase of disease development. Patient committor trajectories visualize how individual conditions evolve over time.

The paper proceeds as follows. The Methods section reviews the optimal RC framework, the committor function, and the free energy profile (FEP). We then present non-parametric RC optimization algorithms, and validation criteria for equilibrium and non-equilibrium sampling, extended to accommodate irregular longitudinal data. Next, we illustrate the framework’s performance on a simple model system under various irregular samplings, demonstrating that approach determines the optimal RC with excellent accuracy. Finally, we assess the sensitivity and robustness of different metrics in validating the optimality of the putative RC across different samplings. We focus on commonly used metrics, such as the AUC, the Mean Squared Error (MSE), and our validation criterion and show that the latter provides the most sensitive and reliable performance. The paper concludes with a discussion.

## 2 Methods

One way to analyze complex multidimensional stochastic dynamics is to project it onto a single coordinate, often referred to as an index, progress variable, order parameter, or reaction coordinate. The dynamics are then described as a diffusion on a free-energy profile (or landscape) as a function of the RC. This provides a simple and intuitive description of the dynamics. While using just one coordinate may result in some loss of information about the original dynamics, if the coordinate is chosen carefully or optimally, the lost information may not be significant. Ideally, by observing the current dynamics of such an optimally selected coordinate, one can predict the future state of the dynamics.

The committor function is known to be an optimal RC for equilibrium dynamics between two boundary states, *A* and *B*. It is defined as the probability of reaching boundary state *B*, starting from point *x*, before reaching state *A* [16, 20]. For example, consider using the committor RC to analyze disease/patient dynamics. Assume longitudinal data, including various blood tests or other biomarkers, has been collected for a relatively large cohort of patients. By applying an optimization algorithm that takes all the patient data as input and computes the committor function as output, this function will predict the probability that a patient will reach state *B* (disease) before reaching state *A* (healthy), which is arguably the most important characteristic. This can be achieved without knowing the exact mechanism of the disease.

However, computing the committor function presents a significant challenge. One common approach in biomolecular dynamics analysis involves approximating optimal RCs through neural networks. This parametric approach requires a specific functional form with numerous parameters, which in turn requires substantial expertise in the system under study. Using too few parameters can lead to poor approximations and suboptimal RCs, whereas an excessive number of parameters increases the risk of overfitting.

To address this challenge and closely approximate the RC, we developed a family of non-parametric RC optimization approaches that do not rely on predefined functional forms with numerous parameters. Initially, a non-parametric method was designed for application to long equilibrium trajectories [21] and subsequently refined [17, 18]. Later, the approach was extended to handle non-equilibrium ensembles of short trajectories saved at constant time intervals [22]. In this paper, we present a further development of this framework to nonequilibrium simulations with imbalanced data and irregular time intervals. This extension enables the realistic analysis of disease progression and patient dynamics using clinical datasets. While detailed descriptions of the methodology and mathematical justification are provided in the referenced works, we offer a brief summary below.

### Cut-based free energy profiles

Consider the reaction coordinate time series *x*(*t*). The cut-based free energy profiles (CFEP) *Z*_*C, α*_ (*x*, Δ*t*) are computed as follows. For every transition in the trajectory from *x*(*t*) to *x*(*t* + Δ*t*), where Δ*t* is the sampling time interval, one adds half the length of the transition raised to the power *α*: |*x*(*t* + Δ*t*) − *x*(*t*)| α*/*2 to all the points between *x*(*t* + Δ*t*) and *x*(*t*) [23, 16].

The CFEP *Z*_*C, α*_ can be used to compute the FEP, the position-dependent diffusion coefficient *D*(*x*), as well as other kinetic properties. In particular, for *α* = −1, the FEP can be calculated as [16]

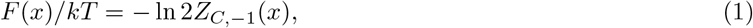

where *k* is the Boltzmann constant and *T* is the temperature. One can calculate the FEP in the conventional way using the probability density of the reaction coordinate in a bin Δ*x*. However, decreasing Δ*x* increases the spatial resolution of the partition function, which also increases the statistical noise. In contrast, for *Z*_*C,−*1_, small Δ*x* does not result in increased noise, allowing for the use of as small Δ*x* as desired. (Jupyter notebooks illustrating the usage of *Z*_*C, α*_ profiles for RC analyses are available at https://github.com/krivovsv/CFEPs.)

### Diffusive model and natural reaction coordinate

Let’s define a putative reaction coordinate time series as *r*(*t*) and the optimal RC, committor, as *q*(*t*). Assume that the optimal RC, the committor *q*(*t*), has been determined. The diffusive model of the dynamics, which allows the exact computation of many important properties of the original dynamics exactly, is defined by the FEP *F* (*q*)*/kT* and diffusion coefficient *D*(*q*) as functions of the committor. The latter is a strongly varying function, as the committor itself is a complex, nonlinear function of the configuration space, which can make it inconvenient for practical applications.

Alternatively, the committor *q* can be transformed or re-scaled to a natural committor 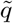, along which the diffusion coefficient is constant. The natural committor 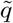 is as optimal as the original committor *q*. Since the diffusion coefficient 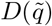 is constant, it is sufficient to show only the FEP 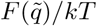 for a complete description of the dynamics [24, 17].

### Non-parametric optimization of the committor

The approaches for determining committors for equilibrium and non-equilibrium samplings are fundamentally similar, differing only in their optimization functionals. A detailed description of the corresponding equations can be found in the Ref. [22].

Here, *X*(*t*) denotes the original multidimensional trajectory times series, i.e., time series of all coordinates that describe patients conditions as a function of time. For simplicity, we denote the time index of the trajectory as *t*. Note that *t* is not a continues variable but rather a set of irregularly sampled time points *t*_*i*_, which, in the case of regular sampling, correspond to *t*_*i*_ = *i*Δ*t*.

### Equilibrium sampling

*Initialization:* A seed RC *r*_0_(*t*) is constructed, which satisfies the boundary constraints. For example, *r*_0_(*t*) = 0 if *X*(*t*) ∈ *A, r*_0_(*t*) = 1 if *X*(*t*) ∈ *B*, and *r*_0_(*t*) = 0.5 otherwise.

*Iterations:* First, a times-series *y*(*t*) is computed from the original trajectory *X*(*t*) (a randomly chosen coordinate or a collective variable). *y*(*t*) is used to improve the current RC *r*(*t*). Specifically, one considers a variation of the RC *δr*(*t*), which can be taken as a small degree polynomial of *y*(*t*) and *r*(*t*) itself

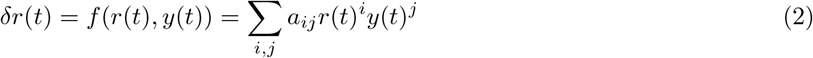

The variation *δr*(*t*) is zeroed on the boundary states to satisfy the boundary conditions. The updated putative RC is given by *r*_*m*+1_(*t*) = *r*_*m*_(*t*) + *δr*_*m*_(*t*), where the index *m* denotes the iteration step. To update the RC, the coefficients *a*_*ij*_ are found by minimizing the following functional [22]

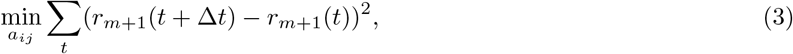

where *r*(*t* + Δ*t*) is the value of the reaction coordinate at the next time step *t* + Δ*t*.

*Stopping:* Iterations stop when the change in the RC time-series during the last *n* iterations, ∥*r*(*t*) − *r*_*−n*_(*t*)∥, is sufficiently small. Different stopping criteria can be specified, such as the number of iterations or the minimal value of the loss function, as an indication of convergence.

### Committor validation criterion for equilibrium sampling

If a putative RC *r* closely approximates the committor *q*, then

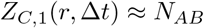

for all *r* and Δ*t*, where *N*_*AB*_ equals to the number of transitions made by the RC time series from boundary state *A* to *B* [16], and *Z*_*C*,1_ is cut profile, which can be directly computed from the time series *r*(*i*Δ*t*), as described earlier. For the committor computed from an equilibrium trajectory, the theoretical lower boundary for the value of the loss function Δ*q*^2^ (Eq. 3) is equal to 2*N*_*AB*_ [16, 17]. Thus, if Δ*r*^2^ ≈ 2*N*_*AB*_, then the RC *r* should closely approximate the committor *q*. This fact can be used as a stopping criteria in the optimization algorithm.

### Non-equilibrium sampling

The algorithm follows the same *Initialization, Iterations* and *Stopping* steps as before. However, for a non-equilibrium ensemble of trajectories, the detailed balance is not satisfied, and thus the committor function does not provide the minimum to the functional (3) any more. Here, we consider a different functional, whose minimum is provided by the committor function [22]

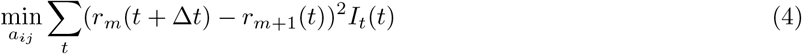

with boundary conditions *r*(*t*) = 0 for *X*(*t*) ∈ *A, r*(*t*) = 1 for *X*(*t*) ∈ *B*. The function *I*_*t*_(*t*) is an indicator function, which equals 1 when *r*_*m*_(*t* + Δ*t*) and *r*_*m*+1_(*t*) belong to the same short trajectory and 0 otherwise. *I*_*t*_(*t*) eliminates all cross-trajectory terms, ensuring that only terms where *r*_*m*_(*t* + Δ*t*) and *r*_*m*+1_(*t*) are from the same trajectory contribute to the sum.

### Committor validation criterion for non-equilibrium sampling *Z*_*q*_

A validation criterion for committor *Z*_*q*_ for non-equilibrium sampling was defined in [22]. While it was successfully applied to the analysis of computer simulated trajectories, the criterion was restricted to relatively small sampling intervals, where the contribution from boundary states was negligible and not taken into account. Here, we generalize the criterion to be applicable for the analysis of patients trajectories, which have non-regular sampling intervals, as well as trajectories of variable length and various boundary conditions.

The derivative *dZ*_*q*_(*x*, Δ*t*)*/dx* can be computed from RC time-series *r*(*i*Δ*t*) as follows [22]

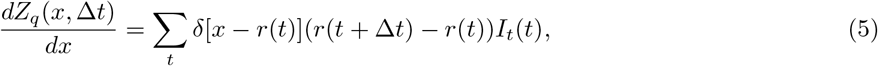

where *x* is a point on the RC, *δ*[·] is the Dirac delta function, *I*_*t*_(*t*) is an indicator function, defined earlier. Thus, *dZ*_*q*_(*x*, Δ*t*)*/dx* at point *x* is proportional to the average displacement ⟨*r*(*t* + Δ*t*) − *r*(*t*)⟨for all segments of trajectories that leave point *x*. For the committor *q*, this average displacement is zero (except at boundary nodes, see below).

*Validation criterion:* If the putative RC closely approximates the committor *q*, then *Z*_*q*_(*x*, Δ*t*) is constant for for all *x* and Δ*t*. Note that, in contrast to the equilibrium case, the constant value here is not informative, as it is defined by the transitions from the state *A* and depends on the sampling.

In practice, *Z*_*q*_(*x*, Δ*t*) is computed by dividing the RC into bins of size Δ*x*. For each time step *t* where *r*(*t*) belongs to bin (*x, x* + Δ*x*), i.e., *x < r*(*t*) *< x* + Δ*x*, the value *dZ*_*q*_(*x*, Δ*t*)*/dx* in bin (*x, x* + Δ*x*) is incremented with *r*(*t* + Δ*t*) − *r*(*t*). Finally, *dZ*_*q*_(*x*, Δ*t*)*/dx* is numerically integrated to obtain *Z*_*q*_(*x*, Δ*t*). *Z*_*q*_(*x*, Δ*t*) deviates from the constant value around the boundaries since the derivative is not zero [22]. These deviations can be eliminated by employing the transition path segment summation scheme [16]. Specifically, one divides all trajectories that visit boundary nodes into segments that either end and/or start with boundary nodes, ensuring no boundary node is within a segment. To compute *Z*_*q*_(*x, k*Δ*t*) for specific value of *k*Δ*t*, each segment ending in a boundary state is extended by *k* steps with the last value. For example, for *k* = 2, a trajectory segment ending in state *B* as […, *r*(*t*_*i−*1_), *r*(*t*_*i*_), *r*(*B*)] is extended for 2 steps as […, *r*(*t*_*i−*1_), *r*(*t*_*i*_), *r*(*B*), *r*(*B*), *r*(*B*)]. This procedure assumes that trajectories are of practically infinite length. For finite-length trajectories, the continued segment should not extend beyond the original trajectory length.

Another modification is required for sampling when a boundary state is also a trap state, i.e., when the trajectory stops at the state, such as when the boundary state corresponds to the death of a patient. In such cases, to keep *Z*_*q*_ constant for the optimal RC (the committor), one must extend the segments up to the length of other trajectories that have not reached the trap state. Assuming the distribution of trajectories length is well described by a Poisson distribution, i.e., the probability of a patient dropping out from the study is approximately constant, the trajectories, that ended up in a trap can be continued with a similar probability to stop, extending the trajectory that ended in a trap with the Poisson distributed length.

To assess whether *Z*_*q*_ deviates from a constant value, one can visually inspect the profiles, or use a descriptive cumulative characteristics, such as maximal standard deviation (*stdev*) of *Z*_*q*_(*k*Δ*t*). This involves computing the standard deviation for each *k* and then determining the maximum. A reasonably small *stdev* indicates that *Z*_*q*_ is approximately constant and therefore corresponds to an optimal RC, whereas a large *stdev* suggests that the coordinate is not optimal.

### Re-weighting factors

For non-equilibrium sampling, re-weighting factors describe the necessary adjustments to the weight of each point to reproduce equilibrium sampling (for an equilibrium trajectory, these factors are equal to 1) [22]. Thus one can obtain an equilibrium FEP along the committor, which can be used to determine important properties of the dynamics exactly. For instance, in the equilibrium free-energy profile *F* (*q*)*/kT*, each computed point *q*(*i*Δ*t*) contributes with the corresponding weight *w*(*i*Δ*t*). For equilibrium *Z*_*C,α*_ cut-profiles, each transition from *q*(*i*Δ*t*) to *q*(*i*Δ*t* + Δ*t*) contributes with the corresponding weight *w*(*i*Δ*t*). For a detailed description of re-weighting factors and the algorithm to compute them, see [22, 19].

### Comparison of predicted (committor) vs observed probabilities

Another way to asses the optimality of the determined RC is to compare the predicted probability of reaching state *B*, which is the determined committor, with the observed probability computed from the trajectories. To do this, the committor RC is divided into bins and, for each bin, the ratio *n*_*A*_*/*(*n*_*A*_ + *n*_*B*_) is computed, where *n*_*A*_ and *n*_*B*_ are the numbers of trajectories reaching states *A* and *B*, respectively. For non-equilibrium sampling, it is important to check that the number of discarded trajectories, which are those that do not reach either of the boundary states, is negligible; otherwise, such an estimate will be biased.

We emphasize that this comparison is different from the standard calibration plot or concordance between predicted and observed probabilities [5]. To illustrate the difference, consider an arbitrary coordinate *r*, such as one describing patient condition. Now, consider the committor as a function of the coordinate *q*(*r*), learned based on training data, which represents the predicted probability. Agreement with the observed probability will show that the model is properly calibrated, and this can be done for any coordinate, even sub-optimal ones.

In our comparison, the predicted probability is the committor function, determined based on the dynamics at the smallest time-scale Δ*t* = Δ*t*_0_, while observed probabilities are determined based on dynamics at largest time-scale, which is the probability of visiting state *B* in the future. Agreement between them is possible only when the putative RC closely approximates the committor and provides a close Markovian approximation to dynamics at all timescales.

## 3 Results

### 3.1 Model system

The described framework of optimal reaction coordinates and the non-parametric approach for their determination have been successfully applied to various model and realistic systems of different complexity, including low- and high-dimensional model systems [21, 22], state-of-the-art atomistic protein-folding simulations [15, 16, 17, 18], chess game [19, 20] among others, illustrating the power and generality of the framework. Here, to clearly demonstrate the fundamental aspects of the approach in analyzing patient dynamics, we consider arguably the simplest model system that has both optimal and sub-optimal RCs. The system consists of two parallel one-dimensional pathways connecting two states, *A* and *B*, described by two variables: 0 ≤ *x* ≤ 1 and *i* = ±1. The variable *x* runs along the pathways, while *i* distinguishes the two pathways. The boundary states are defined as *A* (*x* = 0) and *B* (*x* = 1).

The potential energy *U* (*x, i*) as a function of configuration space is defined

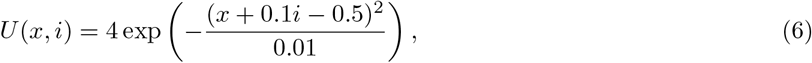

and is shown in Fig. 2a.

**Figure 2:**
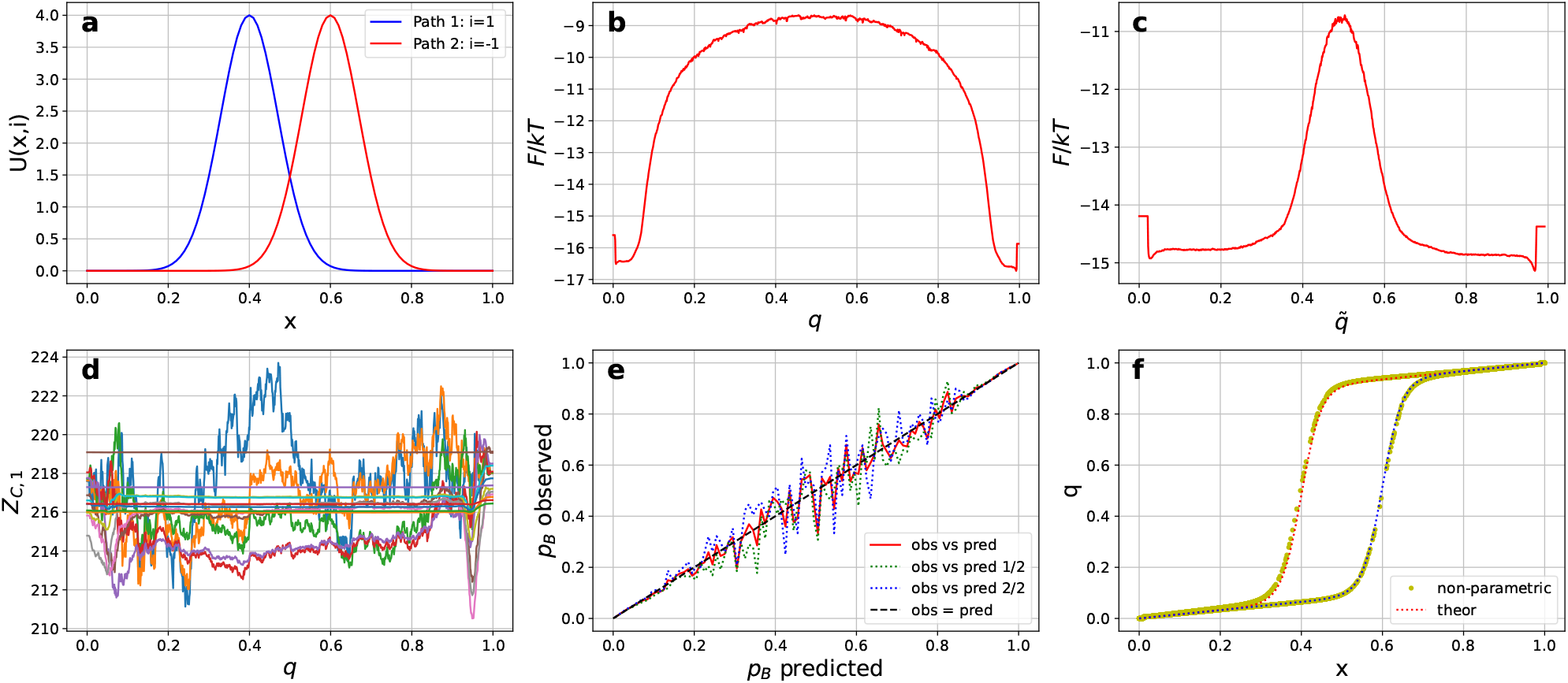
Equilibrium sampling. **a**) Energy landscape with two pathways. The indices *i* = 1 and *i* = −1 correspond to the two distinct pathways; **b**) Free energy as a function of the committor *q* and **c**) natural committor 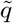, where 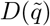 = *const*; **d**) Validation criterion *Z*_*C*,1_(*q, k*Δ*t*0) along the time series *q* for different sampling intervals *k*Δ*t* with *k* = 1, 2, 4…2^16^ remains relatively constant, confirming accuracy of determined committor; **e**) Predicted versus observed pB probabilities: results from the first half of the trajectory (green dotted line), the second half (blue dotted line), and the full trajectory (red line); **f**) Committors computed analytically (red and blue dotted lines) and via the non-parametric approach (yellow line) show excellent agreement.

Stochastic trajectories, modeling the dynamics of patient conditions or disease progression, were generated by simulating diffusive motion within the configuration space of the model system, using a constant diffusion coefficient of *D* = 1, the specified potential energy landscape, and a saving/sampling interval of Δ*t*_0_ = 0.0005. We explore various types of equilibrium and non-equilibrium samplings, ranging from simple to more complex. These include:

#### Equilibrium sampling

A single long equilibrium trajectory with a constant saving interval Δ*t* = Δ*t*_0_, and without traps.

#### Equilibrium sampling

A single long equilibrium trajectory with variable saving intervals Δ*t* = *k*Δ*t*_0_, where *k* = 1, 2, 3, … is a random variable, and without traps.

#### Non-equilibrium sampling

Multiple short trajectories of fixed length with constant Δ*t* = Δ*t*_0_, and without traps.

#### Non-equilibrium sampling

Multiple short trajectories of fixed length with constant Δ*t* = Δ*t*_0_, and a trap in state *B*.

#### Non-equilibrium sampling

Multiple short trajectories of variable length with saving intervals Δ*t* = *k*Δ*t*_0_, where *k* is a random variable, and a trap in state *B*.

The final sampling can be considered a realistic model of patients trajectories derived from clinical datasets, such as EHR. Short non-equilibrium trajectories represent the clinical data of each patient, recorded at irregular time intervals over varying periods. We can combine these short trajectories into a single long trajectory, representing “non-equilibrium sampling with variable Δ*t*”. Two different scenarios can be considered: in the first, multiple trajectories may revisit the boundary states *A* and *B* several times before stopping, corresponding to sampling without traps, where patient trajectories continue to be analyzed even after a disease is diagnosed. In the second scenario, each trajectory terminates as soon as it reaches one of the boundary states, corresponding to sampling with traps, where the trajectory stops once a patient reaches the disease state *B*.

The non-parametric approach has been successfully applied to all sampling types, demonstrating its accuracy and robustness. Below, we present a selection of the most illustrative results. The final outcomes of the analysis in each case are the optimal RC or committor time-series (i.e., the committor value for each frame of every patient’s trajectory) and the equilibrium FEP as a function of the committor. The former enables monitoring of each patient’s current conditions, particular the likelihood of progression to state *B*, such as developing a disease. The latter provides an accurate representation of the overall disease dynamics.

Here, we present the results for equilibrium single trajectory sampled with constant Δ*t* and for the most complex non-equilibrium sampling. The length and number of trajectories were chosen such that the total length of concatenated trajectory is approximately 2000000 steps. Analyses of the three intermediate samplings are provided in the Supplementary information (Figs. S1-S3). Jupyter notebooks for all model analyses are available at https://github.com/krivovsv/D4M.

### 3.2 Equilibrium regular sampling

We first consider equilibrium sampling, which determines the equilibrium FEP and serves as the baseline for comparison with non-equilibrium sampling. Equilibrium sampling represents a hypothetical scenario involving a very long trajectory of a single patient. In contrast, the non-equilibrium sampling discussed later can be interpreted as a method to describe such dynamics using practically available ensembles of short patients trajectories.

A long diffusive equilibrium trajectory was simulated with the following parameters: diffusion coefficient *D* = 1, time step Δ*t*_0_ = 0.0005 and number of steps *N* = 2000000 (the length of the trajectory). To determine the committor *q* we perform non-parametric RC optimization as follows.

#### Initialization

Definition of boundary states: *A* if *x <* 0.01 and *B* if *x >* 0.99. Seed RC: *r*_0_(*t*) = 0 if *x*(*t*) ∈ *A, r*_0_(*t*) = 1 if *x*(*t*) ∈ *B*, and *r*_0_(*t*) = 0.5 otherwise.

#### Iterations

*y*(*t*) is a time-series of either the position *x*(*t*) or the path *i*(*t*), each chosen randomly with equal probability. Polynomials of the 6th degree are used for the variation *δr*(*t*) (Eq. 2); while this specific choice was based on empirical testing, as it consistently ensured stable convergence and reasonable computational cost across all tested scenarios, the results remain robust with respect to the degree selected. To update the RC, the coefficients *a*_*ij*_ are found by minimizing functional (Eq. 3)

#### Stopping

Iterations stop when root mean squared change of the RC after 1000 iterations, being less than 10^*−*4^ 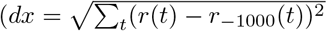, where *r*_*−*1000_(*t*) is the RC calculated at 1000 iterations earlier).

Figure 2 illustrates the results of non-parametric RC optimization applied to the equilibrium sampling. Fig. 2b displays the FEP, *F* (*q*)*/kT*, as a function of the committor *q*. Fig. 2c shows the FEP, 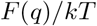, as a function of the re-scaled or natural committor 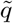, where the diffusion coefficient is constant 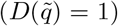. This re-scaling ensures that the stochastic dynamics can be effectively described by the FEP alone [24, 17]. In contrast, using the FEP from Fig. 2b to describe the dynamics is more challenging because the diffusion coefficient *D*(*q*) is a strongly varying function of the committor (see Methods for details). Note that while committor RC *q* always changes from 0 to 1, the natural committor 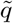 may span a different range, corresponding to dynamics where 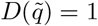. Here, the range of 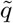 is in agreement with the span of *x*, chosen for the model system, where dynamics is specified with *D*(*x*) = 1.

Figures 2d-2f validate the accurate computation of the committor RC. Fig. 2d shows that the committor time series satisfies the stringent equilibrium committor validation criterion. Specifically, *Z*_*C*,1_(*q, k*Δ*t*_0_) remains relatively constant across all sampling intervals *k*Δ*t*_0_, where *k* = 1, 2, 4…2^16^ represents sampling at every frame, every second frame, every fourth frame, and so on. The variation in ln *Z*_*C*,1_(*q, k*Δ*t*_0_) is bounded within ±0.03, confirming that the RC closely approximates the committor. Furthermore, the number of transitions from state *A* to state *B*, denoted as *N*_*AB*_, directly calculated from the equilibrium trajectory, is 215. This value is close to *Z*_*C*,1_ and the final value of the loss function Δ*r*^2^*/*2 which is 217, further confirming the optimality of the putative RC [16].

Fig. 2e compares the committor probabilities (pB predicted) for each point *x* of the trajectory with the actual observed probabilities (pB observed) of reaching boundary *B* before reaching *A*, calculated directly from the trajectory. The plot demonstrates good agreement between pB predicted and pB observed, with fluctuations around the black dotted line representing the “ideal” observed = predicted plot. To ensure these fluctuations do not exhibit systematic deviations, results are compared for the first half of the trajectory (green dotted line), the second half of the trajectory (blue dotted line), and the full trajectory (red line).

Finally, Fig. 2f shows that the committor as a function of configuration space, *q*(*x, i*) is in excellent agreement with the analytically computed values. For the model system, the committor function can be derived analytically by considering each pathway independently and using the following equation [22]

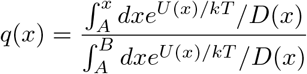

It is important to note that such an analytical computation of the committor is feasible only for such simple model systems. To validate the analysis of real patient data, one must rely on criteria similar to those used in Fig. 2d and 2e.

Fig. 2c presents the FEP describing the overall dynamics of a disease progression. The high barrier, with a height of Δ*F/kT* ~ 4, indicates that the transition from state *A* (healthy) to state *B* (disease) is a rare event. The top of the barrier represents the most critical region for understanding this transition.

One potential strategy for analyzing the etiology of disease progression is to stratify patients based on the values of the optimal coordinate - for example, just before, at the top, and just after the barrier - and then analyze the differences between these groups. The FEP along the optimal coordinate also enables precise computation of key dynamical quantities, such as the mean first passage time (MFPT) and the mean transition path times (MTPT), starting from any position on the coordinate [17, 20].

Here, we consider a model of a disease with two possible pathways, where patient conditions develop in completely different way. In reality, there may be many more pathways of disease development, each with distinct dynamics. Nevertheless, a one-dimensional, single pathway diffusive model of disease dynamics, such as the one shown in Fig. 2c, can be used to compute these properties for all patients trajectories across diverse pathways exactly.

In a clinical context, this approach not only enables the identification of individuals at higher risk of developing specific diseases but also allows for predicting the time frames in which these developments are likely to occur.

### 3.3 Non-equilibrium irregular sampling

In this section, we present the results for the most complex type of non-equilibrium sampling considered, involving multiple short trajectories with variable lengths and saving intervals, Δ*t*, as well as a single trap at boundary *B*. This type of sampling serves as a realistic model for analyzing disease or patient dynamics based on clinical datasets. The trap in the disease state corresponds to discarding the portion of the trajectory after a patient reaches the disease state and begins treatment, as this phase is not captured in the clinical data. Excluding this segment ensures that the analysis remains uncontaminated by inconsistent information.

A total of 220000 short trajectories are simulated with the following parameters: diffusion coefficient *D* = 1, maximum number of steps per trajectory *nsteps* = 200, and an initial time step Δ*t*_0_ = 0.0005. Each trajectory has a 0.05 probability of termination at every step; it also terminates upon reaching boundary *B*. To introduce a variable time step, trajectory coordinates were randomly saved at each time step with a probability of 0.5. Consequently, the lengths of the trajectories ranged from 1 to 200, with an average of 10 steps. This setup reflects the nature of clinical data, where patient records are irregular and typically consist of a few data points on average. The short trajectories are concatenated into a single long trajectory, comprising in a total of ≈ 2000000 steps.

To determine the putative time-series *r*(*t*), which closely approximates the committor *q*(*t*) we perform the non-parametric RC optimization. The initialization, choice of *y*(*t*) and stopping criterion are consistent with those used in equilibrium optimization. Polynomials of the 6th degree are employed for the variation *δr*(*t*) (Eq. 2). The RC is updated by finding the coefficients *a*_*ij*_ through minimization of the functional (Eq. 4).

Figure 3 demonstrates that the non-parametric approach can accurately determine the committor time series and the corresponding equilibrium FEP from non-equilibrium sampling, representing realistic clinical data. This approach effectively addresses irregular saving or observation intervals. Fig. 3a shows that the committor time series satisfies the stringent non-equilibrium committor validation criterion. Specifically, *Z*_*q*_(*q, k*Δ*t*_0_) remains relatively constant across all sampling intervals *k*Δ*t*_0_, where *k* = 1, 2, 4 2^16^. Deviations are within the range of statistical fluctuations ( ±3%), consistent with *Z*_*q*_(*q*, Δ*t*_0_), which by construction should remain constant except for such fluctuations.

**Figure 3:**
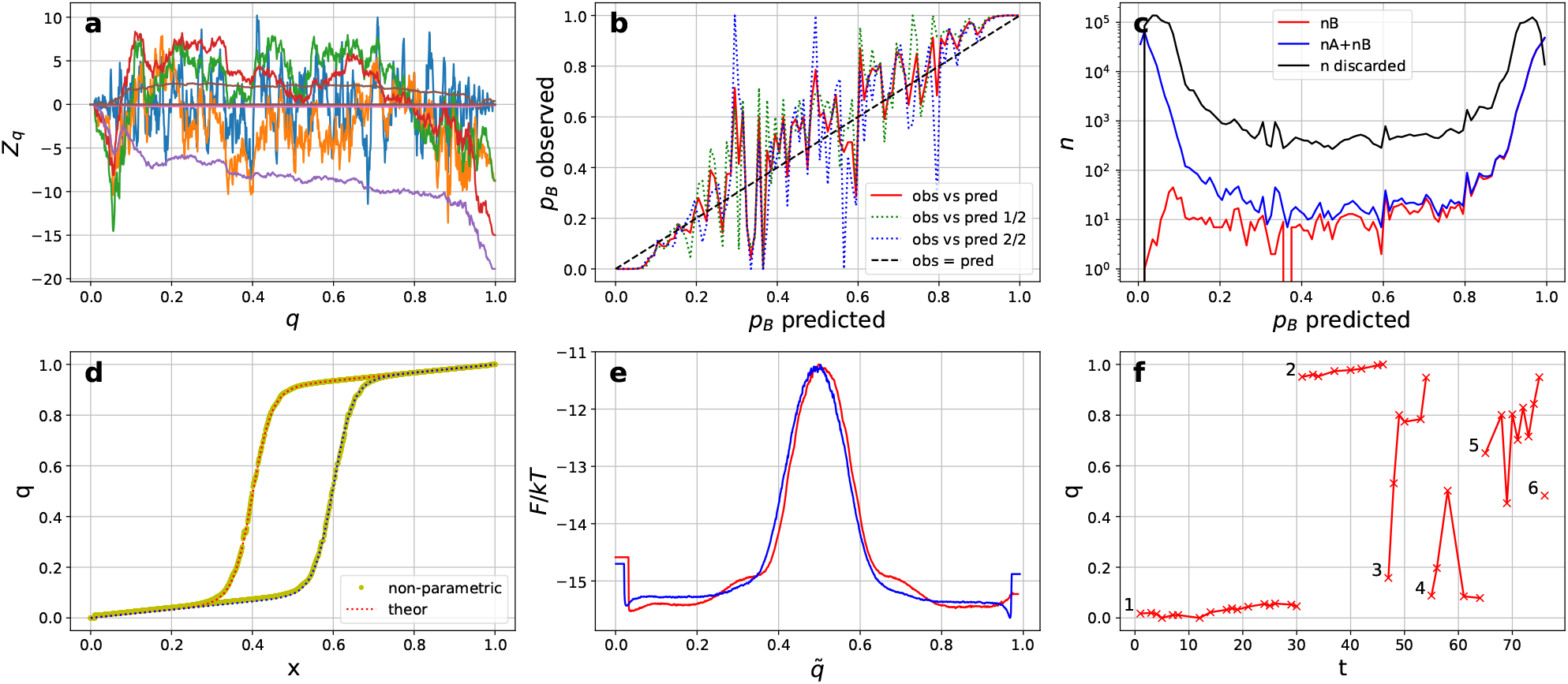
Non-equilibrium irregular sampling. **a**) Validation criterion: *Z*_*q*_ (*q, k*Δ*t*) along the time series *q* for *k* = 1, 2, 4..2^16^ remains relatively constant, confirming accurate committor computation; **b**) Predicted versus observed pB probabilities: results from the first half of the trajectory (green dotted line), the second half (blue dotted line), and the full trajectory (red line); **c**) The total observed number of trajectories passing through *q* that have reached either *A* or *B* (blue line), that have reached state *B* (red line), and that have not reached boundaries (black line); **d**) Committors computed analytically (red and blue dotted lines) and via the non-parametric approach (yellow line) show excellent agreement; **e**) FEPs along the natural committor 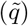 calculated for non-equilibrium sampling (red line) and compared with the FEP computed for equilibrium sampling (blue line); **f**) Selected committor trajectories illustrating how “patient’s” conditions could develop over time.

Fig. 3b compares the predicted and observed pB probabilities, revealing significant deviations from the middle line. Systematic discrepancies are evident near the boundaries: the red curve (observed vs. predicted) dips below the ideal line (black dotted line) at (*q* ~ 0) and rises above it at (*q* ~ 1). These deviations occur because the observed probabilities cannot be accurately estimated, as many trajectories fail to reach the boundary states due to their relatively short lengths. The substantial number of such discarded trajectories is illustrated in Fig. 3c (black line).

Fig. 3d demonstrates that the committors computed analytically and via the non-parametric approach from the non-equilibrium sampling are in excellent agreement.

A non-equilibrium FEP along the committor (not shown) has no significant meaning, as it is non-equilibrium and dependent on sampling. However, re-weighting factors can be applied to compute the equilibrium FEP (see Methods for details). Fig. 3e illustrates the equilibrium FEP (after re-weighting) along the natural coordinate 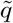 (red line). The results show excellent agreement with the FEP obtained from equilibrium sampling (blue line) (see Fig. 2c). This confirms that it is possible to determine the equilibrium FEP from non-equilibrium sampling, such as clinical data, providing an accurate description of the overall disease dynamics.

Fig. 3f shows several short committor trajectories, illustrating different patterns of how “patients” conditions could develop over time, i.e., how each patient’s individual likelihood of disease onset changes dynamically. Patient 1 stays in the healthy basin for the entire duration of their trajectory; while patient 2’s trajectory belongs to the disease basin and reaches state *B* (disease) at the final step; patient 3 starts in the healthy basin, transitions to the disease basin, and ends with a high likelihood of disease onset; while trajectory of patient 4 starts in the healthy basin, develops to the transition state and then returns to the healthy basin, illustrating the stochastic nature of the dynamics; patient 5 starts in the disease basin, visits the transitions state, however returns to the disease basin and ends with a high likelihood of developing the disease; trajectory of patient 6 consists of a single point. While most trajectories do not reach the boundaries, their ultimate outcomes can still be predicted, a task that would be impossible without considering dynamics of disease, which is not feasible in simple classification models based on case/control design. These trajectories, along with the representation of the disease as a landscape that highlights the critical transition region (Fig. 3e), provide a visual and intuitive way for the proposed framework to explains the analysis results. This representation makes it easier to explain aspects of complex disease dynamics to clinicians and patients, enabling clearer insights into disease progression and risks.

### 3.4 Metrics Analysis

In the previous section, we have demonstrated that the non-parametric approach can accurately compute the committor and the corresponding FEP from various types of equilibrium and non-equilibrium sampling. In this section, we evaluate the performance of various metrics that can be used to differentiate between optimal and sub-optimal coordinates, denoted as *q* (optimal) and *q*_*sub*_ (sub-optimal). Specifically, we focus on three metrics: the AUC, the MSE, and the optimality criterion *Z*_*q*_ introduced earlier. The AUC metric is widely used to evaluate the performance of binary classification models, while the MSE quantifies the average error between predicted and actual values. Lastly, *Z*_*q*_ serves as an indicator of RC optimality.

We present results for three systems: (1) an “ideal system”, specifically chosen to highlight cases where all three metrics perform well and distinguish between optimal and sub-optimal RCs; (2) a “typical system”, as used in the previous sections; and (3) a system with imbalanced data. These results emphasize the robustness of the *Z*_*q*_ metric, which we propose as a general metric for the analysis of disease dynamics, addressing recent concerns regarding the limitations of the predominantly used AUC metric [1, 12].

We begin with an “ideal system” where all three metrics demonstrate strong performance. The potential energy function *U*_1_(*x, i*) differs slightly from that shown on Fig. 2a

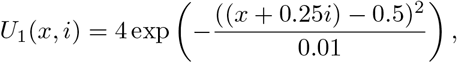

with barriers located at *x* = 0.25 and *x* = 0.75. This spacing places the two barriers farther apart compared to the previous model.

A total of 10000 relatively short trajectories, each 2000 steps in length, were simulated using a diffusion coefficient of *D* = 1 and a constant time step of Δ*t*_0_ = 0.0005. The system includes two traps, meaning that each trajectory terminates upon reaching one of the boundaries, *A* or *B*. A relatively high number of steps ensures that the majority of trajectories reach one of the boundaries, enabling reliable performance of the MSE metric and accurate estimation of the observed pB, since the number of discarded trajectories is negligible. These short trajectories were then concatenated into a single long trajectory, consisting of approximately 2,000,000 steps in total.

Next, we construct the optimal RC, the committor *q*, and the sub-optimal coordinate *q*_*sub*_, and compare the performance of different metrics in evaluating these two coordinates. For both *q* and *q*_*sub*_, we employ a non-parametric approach.

The sub-optimal coordinate *q*_*sub*_(*x*) is derived by using only *x*(*t*) as the collective variable during iterations of non-parametric optimization. In this case, the “committor” is constructed as a function of *x* alone, rather than the entire configuration space consisting of both *x* and *i*. Since the information from the *i* time-series is missing, the resulting coordinate *q*_*sub*_(*x*) is sub-optimal.

Figure 4 presents a comparison of metrics for the “ideal” system, evaluating the sub-optimal coordinate *q*_*sub*_ and the committor *q*. Figs. 4a and 4d display the performance of the validation criterion *Z*_*q*_, demonstrating that *Z*_*q*_(*q*_*sub*_, *k*Δ*t*_0_) is not constant across different sampling intervals *k*Δ*t*_0_, whereas *Z*_*q*_(*q, k*Δ*t*_0_) remains relatively constant for the committor *q*. The calculated maximum standard deviation (*stdev*) of *Z*_*q*_ is notably large at 67.5 for *q*_*sub*_, but reasonably small at 3.55 for *q*.

**Figure 4:**
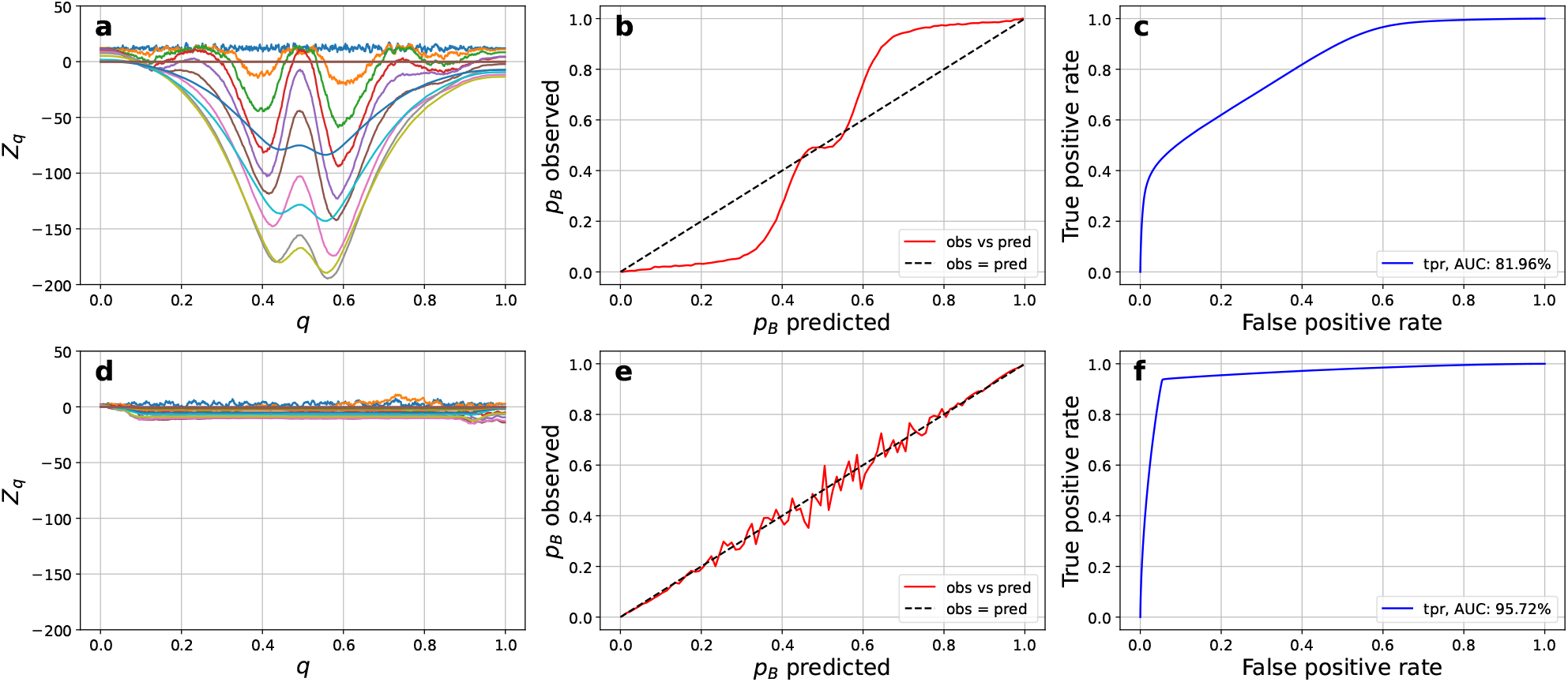
Comparison of metrics for the “ideal” system. The first row (panels **a, b, c**) presents the results for the sub-optimal RC, *q*_*sub*_, while the second row (panels **d, e, f**) shows the results for the optimal committor, *q*. **a, d**) Validation criterion: *Z*_*q*_ (, *k*Δ*t*_0_) for *k* = 1, 2, 4, …, 2^16^; **b, e**) Predicted versus observed pB probabilities; **c, f**) Truepositive rate (ROC curve, blue line) and the threshold (red line) as functions of the false positive rate.

Figs. 4b and 4e show the predicted versus observed pB probabilities. These plots reveal a strong agreement for *q*, but not for *q*_*sub*_.

The AUC metric, shown in Figs. 4f and 4c, highlights a high discrimination power of 95.7% for *q*, compared to 81.96% for *q*_*sub*_. Additionally, the MSE for *q*_*sub*_ is 0.18, while for *q* it is significantly lower at 0.05.

In the case of a carefully designed “ideal” system, where all metrics are expected to perform well, the metrics effectively distinguish between the sub-optimal coordinate *q*_*sub*_ and the optimal committor *q*, consistently demonstrating higher accuracy and better performance for the committor.

Next, we consider the model described in Section 3.2, which represents a more “typical” scenario that closely resembles realistic clinical patients trajectories. In practice, such trajectories are often short, consisting of only a few time points recorded at irregular intervals. Additionally, most trajectories do not reach a boundary state within the observed time-frame.

Figure 5 compares metrics for the “typical” system. Figs. 5a and 5d display the optimality criterion *Z*_*q*_, revealing that *Z*_*q*_(*q*_*sub*_, *k*Δ*t*_0_) varies significantly across sampling intervals *k*Δ*t*_0_, while *Z*_*q*_(*q, k*Δ*t*_0_) remains relatively constant for *q*. Notably, the *stdev* of *Z*_*q*_ is 113.8 for *q*_*sub*_ but only 6.94 for *q*, emphasizing the latter’s constancy.

**Figure 5:**
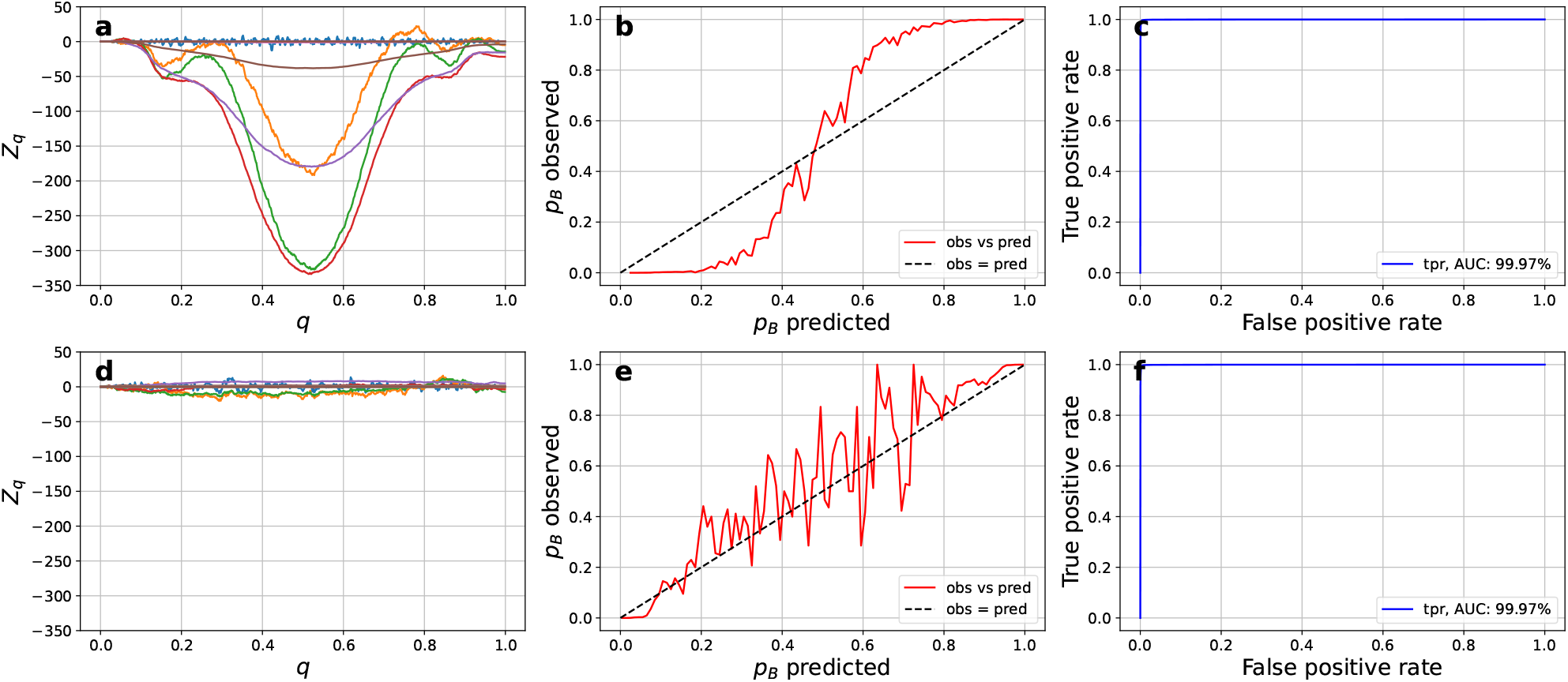
Comparison of metrics for the “typical” system. Notations are the same as in Fig. 4.

Figs. 5b and 5e illustrate predicted versus observed pB probabilities. These probabilities show no clear agreement for either *q*_*sub*_ or *q*. For *q*, the probabilities fluctuate around the ideal (black dotted) line but with substantial deviations. In the “typical” model, many trajectories fail to reach the boundary states due to their short lengths, leading to a considerable number of discarded trajectories, as shown in Fig. 3c. Consequently, observed probabilities cannot be accurately estimated. This contrasts with the “ideal” model, where trajectories were explicitly generated to reach boundary states.

The AUC metric, presented in Figs. 5c and 5f, does not differentiate between the two coordinates at all, showing high discrimination power (99.97%) for both *q*_*sub*_ and *q*. Similarly, the MSE is low for both coordinates, with values of 0.0015 for *q*_*sub*_ and 0.0006 for *q*.

Lastly, we examine a model system representative of imbalanced data, where the population of the disease basin is significantly smaller than that of the healthy basin, reflecting a more realistic scenario. This system is defined by a potential energy function composed of two sigmoid terms:

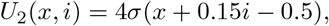

where the right minimum is effectively non-existent. Here, the sigmoid function is given by *σ*(*s*) = 1*/*(1+exp^*−s*^).

A total of 115000 short trajectories are simulated using the following parameters: diffusion coefficient *D* = 1, number of steps per trajectory *nsteps* = 200, and an initial time step Δ*t*_0_ = 0.0005. Each trajectory has a termination probability of 0.025 per step or ends upon reaching boundary *B*. To introduce a variable time step, trajectory coordinates were randomly saved at each step with a probability of 0.5. The short trajectories are concatenated into a single long trajectory, comprising in a total of approximately 2000000 steps.

Figure 6 compares metrics for the “imbalanced data” system. The results are similar to those observed in the “typical” model system. The optimality criterion (Figs. 6a and 6d) reveals that *Z*_*q*_(*q*_*sub*_, *k*Δ*t*_0_) exhibits significant variability, while *Z*_*q*_(*q, k*Δ*t*_0_) remains relatively constant for *q*. The *stdev* of *Z*_*q*_ is 50.12 for *q*_*sub*_ but only 6.15 for *q*, emphasizing the latter’s constancy.

**Figure 6:**
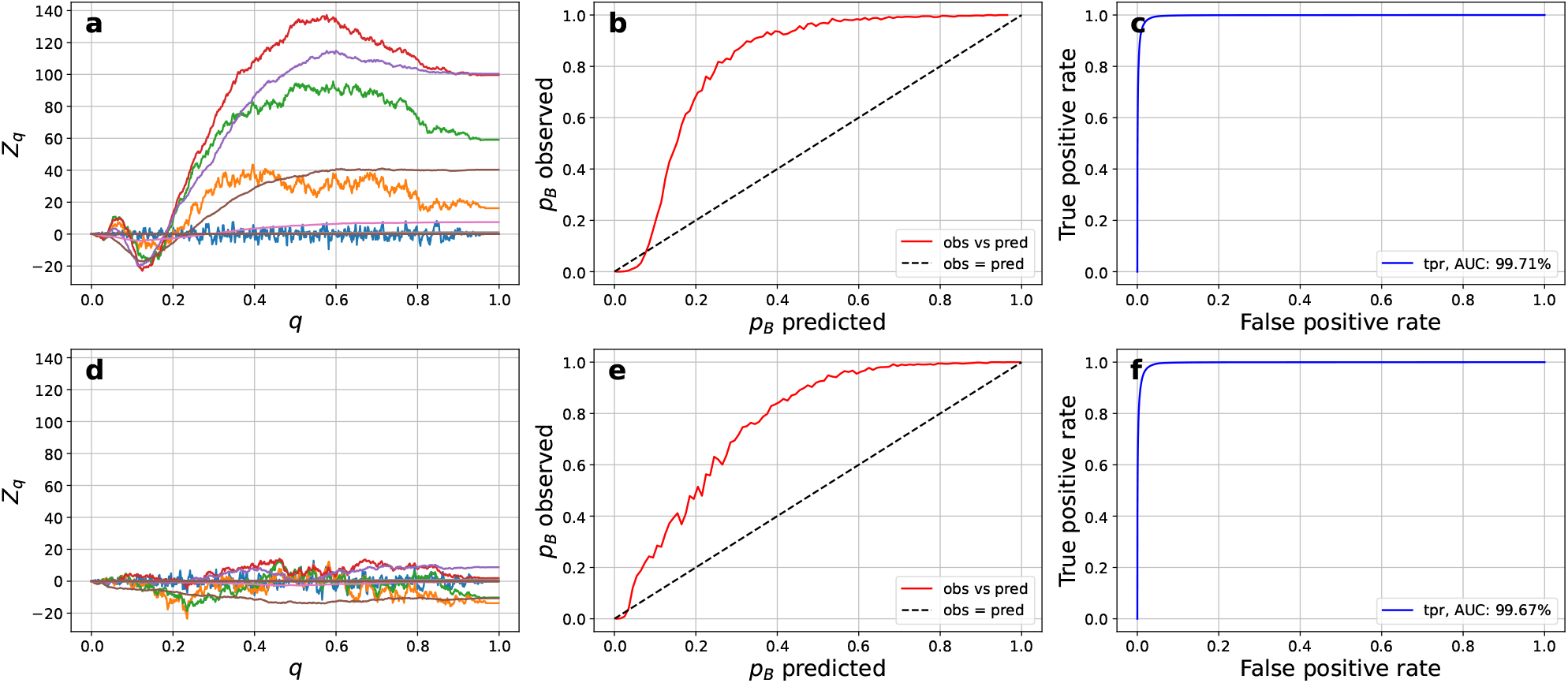
Comparison of metrics for the “imbalanced data” system. Notations are the same as in Fig. 4.

Figs. 6b and 6e illustrate predicted versus observed pB probabilities of reaching the disease basin, showing no agreement for either *q*_*sub*_ or *q*. As before, this disagreement arises from the short length of the trajectories, as most fail to reach the boundary states. Consequently, the observed probabilities cannot be reliably estimated.

The AUC metric, shown in Figs. 6c and 6f, does not differentiate between the two coordinates, showing high discrimination power (99.7%) for both *q*_*sub*_ and *q*. Similarly, the MSE is low for both coordinates, with values of 0.01 for *q*_*sub*_ and 0.008 for *q*.

Additionally, we evaluated the performance of precision-recall curves, recommended for skewed datasets [13], as well as Kaplan-Meier (KM) curves and time-dependent AUCs commonly used in survival analysis, for distinguishing between optimal and sub-optimal RCs (see Supplementary Information, Figs. S4-S6), and obtained similar conclusions.

In summary, the results demonstrate, that unless the model system is carefully designed, the performance of standard metrics, such as AUC, MSE, KM curves, or predicted vs observed probabilities in differentiating between optimal and sub-optimal RCs for describing dynamics is rather weak. In contrast, *Z*_*q*_ validation criterion consistently showed robust and strong performance across all three systems considered. One reason for the generally poor performance of metrics like AUC is their cumulative nature. These metrics aggregate contributions from all regions of the optimal RC. Hence, the transition region, being the most critical for describing disease dynamics, contributes negligibly compared to other regions due to its small population as the dynamics are a rare event.

## 4 Concluding Discussion

Traditional static disease analysis methodologies are inherently limited in their ability to extract dynamic information from complex longitudinal datasets. In contrast, models of disease dynamics offer a fundamentally superior approach by modeling disease progression as a stochastic process, potentially capturing the full informational content embedded within these datasets. Ideally, such models should describe the entire trajectory of disease progression, from healthy to diseased states, with particular emphasis on the critical transition phase, which often holds the key to understanding disease onset.

Finite state Markov chains offer the most comprehensive description of disease dynamics. They serve as a valuable theoretical framework, highlighting the potential capabilities of such models for disease analysis. However, the practical implementation of this brute-force approach towards dynamics modeling faces significant challenges, such as the exponential growth of the configuration space and the correspondingly massive data requirements for accurate training.

To address these limitations, we propose an alternative, practically feasible approach for constructing a foundation model of disease dynamics. Instead of using discrete Markov chains, our method describes disease progression as a diffusion process on a free energy landscape, parameterized by optimal reaction coordinates. This approach preserves the fundamental advantages of dynamical modeling while overcoming the computational and data limitations of the brute-force method.

Our methodology has been developed and rigorously tested on state-of-the-art atomistic protein folding trajectories [16, 17, 18, 22], as well as on various longitudinal data [9, 19]. In this work, we specifically highlight the method’s effectiveness in analyzing disease dynamics, particularly its ability to handle highly irregular longitudinal datasets – a persistent challenge for conventional ML and AI approaches. Additionally, we introduce a new validation criterion for validating optimal coordinates in clinical datasets. This criterion demonstrates remarkably robust and sensitive performance, surpassing standard metrics such as AUC and MSE, especially in scenarios involving imbalanced data. By focusing on the critical transition regions that are most relevant for understanding disease dynamics, our method overcomes the inherent weaknesses of cumulative metrics like AUC, which tend to dilute the contributions of these key regions. Note also that sensitivity or the power of the criterion to distinguish between optimal and sub-optimal coordinates grows with the size of the dataset (*N*), as systematic deviations for the former scale as ~ *N*, while statistical fluctuations for the latter scale as 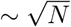.

Another fundamental advantage of the validation criterion over standard metrics such as AUC and MSE is that these traditional metrics lack a theoretical upper bound beyond which improvement is inherently impossible. In contrast, if the validation criterion confirms that a RC derived from the current set of input variables (features) is optimal, then further expanding the input set will not enhance the RC’s ability to describe the disease. Consequently, predictive metrics like AUC will also not improve, even if their current values are relatively low. The analysis determines an equilibrium free energy landscape as a function of the optimal reaction coordinate, mapping each patient’s position within this landscape. This enables a deeper understanding of a patient’s current condition and allows for predicting future developments, such as the likelihood of disease onset. The landscape provides an intuitive, yet qualitatively accurate visualization of disease dynamics. For diseases that are rare events - such as the model system considered here - the landscape features a narrow, high barrier separating two basins, representing the critical transition from a healthy to a diseased state. It can be used for patient stratification, facilitating more detailed analyses of this transition or guiding targeted treatments with drugs designed for specific phases of disease progression.

For conditions that develop gradually over time, such as Alzheimer’s disease or aging, the free energy landscape is expected to exhibit a generally downhill profile, with disease progression modeled as downhill diffusion towards the disease. In such cases, a continuously varying optimal coordinate offers a more refined and accurate description compared to coarse-grained models based on discrete thresholds [7]. Furthermore, constructing free energy landscapes as functions of multiple optimal reaction coordinates enables the identification of metastable states and distinct disease pathways [18], which can be used for defining disease states and different disease development pathways based on patient/disease dynamics in an objective, data-driven way.

A key limitation of the proposed framework is its reliance on longitudinal data to fully demonstrate its power. While the optimization procedure can, in principle, operate with just two time points, the accuracy of the validation criterion, which evaluates the optimality of dynamic descriptions across different time scales, depends heavily on the length of the trajectories. Longer trajectories allow probing longer timescales and improve reliability of the criterion. Although such longitudinal data are more difficult and costly to obtain, their availability is steadily increasing.

It is instructive to compare our approach with state-of-the-art methods for analyzing longitudinal patient data. Since few existing approaches explicitly focus on disease dynamics, we broaden the comparison to related frameworks that align with this central idea. Hidden Markov models (HMMs), for instance, have been used for prognostic disease prediction [10]. As discussed extensively earlier, while Markov models are arguably the most informative construct for the description of dynamics, they have a number of significant shortcomings, addressed by our framework.

Survival analysis (SA), or time-to-event analysis, comprises a range of methods for estimating the distribution of time until an event, such as disease onset or death [8, 7, 25, 26, 27]. By considering time-to-event as a reaction coordinate, analogous to the mean first passage time reaction coordinate [22], SA and our framework can be directly related. One of the most attractive features of SA is its ability to incorporate partially censored and truncated data in the analysis. Our approach, however, explicitly models the underlying dynamics, offering greater flexibility for handling irregular datasets, as demonstrated here. To illustrate the difference, consider slowly progressing conditions such as Alzheimer’s or Parkinson’s. Instead of following a single cohort for decades until enough events occur, our framework combines many short patient trajectories (e.g., five-year records) from individuals at different stages to reconstruct a coherent long-term picture of disease dynamics. This enables estimation of long-term outcomes without extended follow-up. Once built, the model can provide survival probabilities, time-to-event estimates, and other prognostic measures for any stage, even when most data remain censored and hazard estimation is unreliable. In this sense, the framework generalizes SA by enabling long-term risk assessment from short-term observations.

Molecular aging clocks use machine learning models trained on omics data to estimate chronological and biological age and assess the effectiveness of anti-aging or disease-preventive interventions [28, 29, 30]. Most approaches rely on non-longitudinal datasets and fit (non-)linear regression models to chronological age, despite its limitations as a measure of aging progress. Given the complex stochastic nature of aging dynamics, individuals with identical chronological ages may exhibit significant variability in their development trajectories [29]. Consider time from birth (age) and time to death as variables measuring the overall progress of organism development. By using longitudinal datasets, our framework allows for the determination of optimal coordinates, offering a rigorous, data-driven description of aging dynamics. An interesting practical questions is how accurately can one approximate the optimal coordinate based on practically available data and how accurately one can predict the likelihood of death using such a coordinate.

Inspired by the success of large language models (LLM) [31, 32, 33], a new family of approaches has recently emerged. These approaches, often referred to as foundation models, use transformers (or their extensions) to model disease dynamics from longitudinal clinical or biomedical data [3, 34, 35, 36, 37, 38, 39]. These largescale AI models are trained on vast, diverse, typically unlabeled data, applicable to a wide range of scenarios. The major advantage of foundation models is they can serve as a basis for downstream applications and can be fine-tuned for specific tasks with significantly less labelled training data than standard AI models. The latter is of key importance in healthcare applications, where accurate disease identification is often resource-intensive and costly. For example, BrainLM, a foundation model trained on large fMRI datasets, captures brain activity dynamics and can be used to study disease progression [40]. It reconstructs masked brain activity sequences, including previously unseen data, and can be fine-tuned to predict clinical variables such as age, anxiety, and PTSD, as well as forecast future brain states. Roughly speaking, the transformers are used to parameterize the transition probability matrix in the discrete Markov chain approach. It is of interest how well such advanced neural network architectures can mitigating the shortcomings of the brute-force Markov chain approach in the description of disease dynamics. In particular, state-of-the-art LLM use hundreds of billions of parameters and require trillions of data points for training. In contrast, the configuration space of disease dynamics is vastly larger, while the available data is considerably more limited. An important question is how to mitigate the tendency to hallucinate, a well documented behavior of LLM [41]. An interesting application would be using the representations of disease dynamics, learned by a foundation model, to determine optimal coordinates for clinical and disease modeling. An important question is whether such models, trained predominantly on healthy dynamics, contain enough information to accurately describe the much rarer dynamics of disease, i.e., whether they can be used to obtain accurate optimal coordinate, especially in the most critical, transition region. LLMs also provide an illustrative parallel of the power of such dynamical approaches for the analysis of diseases. Just as LLMs generate valuable responses by learning language dynamics without, arguably, true semantic understanding, a model capturing disease dynamics could provide meaningful insights without complete mechanistic understanding [34]. These possibilities motivate the development of our framework as a practically feasible approach to constructing foundation models for disease dynamics.

The framework proposed here can employ an optimal reaction coordinate parameterized with a neural network featuring advanced architectures, such as recurrent neural networks or transformers [35, 36]. However, selecting an appropriate functional that is sufficiently flexible to allow for accurate approximation of the optimal reaction coordinate requires extensive expertise, particularly given the diverse multi-modal data structures used for storing patient information across many hospitals. In contrast, the non-parametric approach offers greater flexibility and does not require extensive expertise with the system or network architectures. It can be considered a robust solution for identifying the optimal coordinate using the best possible parameterization. This approach can serve as an easily deployable baseline model, providing an initial indication of the achievable accuracy from the data. If the model shows promising results, it can be followed by the selection and adoption of an appropriate neural network architecture.

Alternatively, the proposed framework of optimal reaction coordinate, combined with the non-parametric approach and the validation criterion, suggests a new and intriguing workflow, different from the conventional one, where an ML/AI model is trained on one dataset (e.g., from one hospital), validated on another (e.g., from other hospitals) and then deployed to be used on other datasets. This conventional workflow implicitly assumes that all hospitals have compatible EHR systems and that the other datasets are not out of distribution (OOD) with respect to the training dataset, which is often not true in practice [4, 42]. The proposed framework allows for the development and validation of models on the same dataset by leveraging the statistical independence of *Z*_*q*_(*x*, Δ*t*) for different Δ*t*, thereby eliminating the OOD problem. Specifically, *Z*_*q*_(*x*, Δ*t* = Δ*t*_0_) = const does not automatically imply that *Z*_*q*_(*x*, Δ*t*) = const for other Δ*t*. This is only true when the putative reaction coordinate closely approximates the committor. The non-parametric approach optimizes the coordinate for Δ*t* = Δ*t*_0_, leading to *Z*_*q*_(*x*, Δ*t*_0_) = const, while the validation criterion tests *Z*_*q*_ for all Δ*t*. Together with the non-parametric approach, this framework allows for the model to be re-trained from scratch for every new batch of patients, thereby also eliminating the issue of calibration drift. Calibration drift occurs when a model’s predictions become less accurate over time due to changes in population demographics, disease prevalence, clinical practice, and the healthcare system [6, 42].

Although the development of dynamical models for disease progression is still in its early stages, these models hold great promise for enabling a more comprehensive analysis of longitudinal datasets and disease characterization. The optimal RC framework presented here offers one possible methodological pathway toward realizing this potential by shifting our understanding of disease description from static snapshots to dynamic, probabilistic trajectories. It provides a theoretically optimal representation of stochastic disease dynamics. While this method has been thoroughly validated in other areas, such as the complex case of protein folding dynamics, it still needs to be validated in clinical settings. The developed methodology is currently being applied to analyze the onset of acute kidney injury using longitudinal clinical data. Preliminary results demonstrating the feasibility of the approach are reported in [43]

## Data availability

The Python library implementing the framework described in this study is publicly available at https://github.com/krivovsv/optimalrcs. Analysis notebooks are provided in the accompanying Jupyter notebooks at https://github.com/krivovsv/D4M Additional information is available from the corresponding author on reasonable request.

## Acknowledgments

The author gratefully acknowledge the support from Kidney Research UK (KRUK) award number (RP 018 20230628).

## Author contributions

P.B. designed the models, performed the simulations and data analysis, and drafted the manuscript. S.K. developed the methodology, supervised the work, and edited the final manuscript.

## Additional information

### Supplementary Information

