## Supplementary Information for "Data Driven Disease Dynamics Models"

P. Banushkina[1], S. Krivov[1,2,*]
[1]Faculty of Biological Sciences, University of Leeds,
Leeds LS2 9JT, United Kingdom
[2]Astbury Center for Structural Molecular Biology, University of Leeds,
Leeds LS2 9JT, United Kingdom
*

August 3, 2026

### Equilibrium sampling: single long trajectory with variable saving intervals $\Delta t$

A single long trajectory of length $N = 4000000$ was simulated with the following parameters: diffusion coefficient $D = 1$ and initial time step $\Delta t_0 = 0.0005$. To introduce a variable time step, trajectory coordinates were randomly saved at each time step with a probability of 0.5. Supplementary Fig. S1 shows that the approach accurately computes the committor time-series.

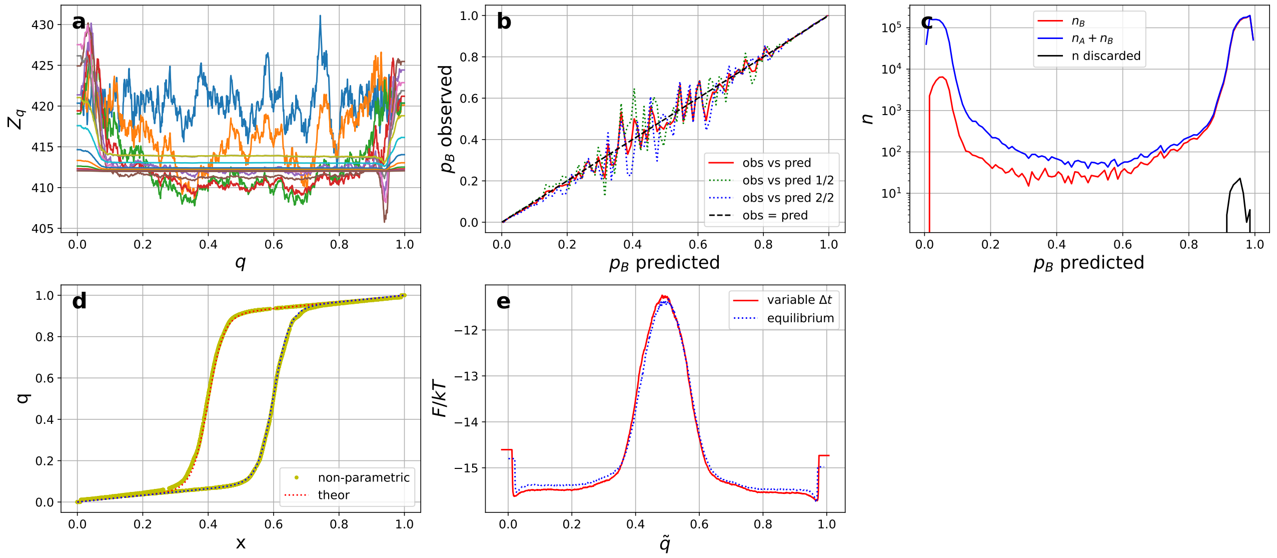


Supplementary Figure S1: Equilibrium sampling with variable sampling interval $\Delta t$. **a**) Validation criterion: $Z_q(q, k\Delta t)$ along the time series $q$ for $k = 1, 2, 4, .., 2^{16}$ remains relatively constant ($\pm 3\%$), confirming accurate committor computation; **b**) Predicted versus observed pB probabilities: results from the first half of the trajectory (green dotted line), the second half (blue dotted line), and the full trajectory (red line); **c**) The total observed number of trajectories passing through $q$ that have reached either $A$ or $B$ (blue line), that have reached state $B$ (red line), and that have not reached the boundaries (black line); **d**) Committors computed analytically (red and blue dotted lines) and via the non-parametric approach (yellow line) show excellent agreement; **e**) FEPs along the natural committor ($\tilde{q}$), calculated for sampling with variable (red line) and constant (blue line) sampling interval $\Delta t$.

### Non-equilibrium sampling with multiple short trajectories of fixed length

A total of 20000 short trajectories were simulated with the following parameters: diffusion coefficient $D = 1$, maximum number of steps per trajectory $nsteps = 100$, and time step $\Delta t_0 = 0.0005$. The short trajectories were concatenated into a single long trajectory, comprising a total of 2000000 steps. Supplementary Fig. S2 shows that the approach accurately computes the committor time-series. Free energy as a function of the committor

RC, computed directly from the time-series (green dashed line in Supplementary Fig. S2e) differs from that obtained for the equilibrium sampling (Fig. 2c or blue dotted line in Supplementary Fig. S2e), confirming that the sampling is non-equilibrium. The equilibrium FEP, which describes the stochastic dynamics, can be recovered by re-weighting the non-equilibrium trajectories (see Methods), and is shown by the red line in Supplementary Fig. S2e.

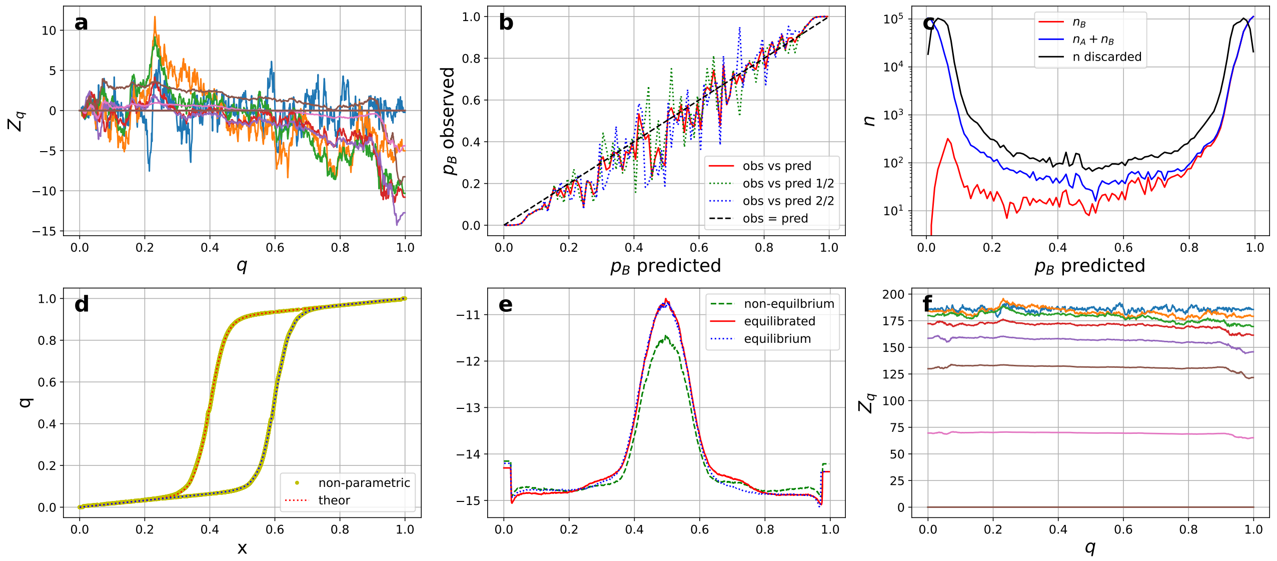


Supplementary Figure S2: Non-equilibrium sampling with multiple short trajectories of fixed length. **a**) Validation criterion: $Z_q(q, k\Delta t)$ along the time series $q$ for $k = 1, 2, 4, .., 2^{16}$ remains relatively constant ($\pm 3\%$), confirming accurate committor computation; $Z_q$ values are shifted vertically for ease of comparison; panel **f**) shows the unshifted curves; **b**) Predicted versus observed pB probabilities: results from the first half of the trajectory (green dotted line), the second half (blue dotted line), and the full trajectory (red line); **c**) The total observed number of trajectories passing through $q$ that have reached either $A$ or $B$ (blue line), that have reached state $B$ (red line), and that have not reached the boundaries (black line); **d**) Committors computed analytically (red and blue dotted lines) and via the non-parametric approach (yellow line) show excellent agreement; **e**) FEPs along the natural committor ($\tilde{q}$), calculated for non-equilibrium sampling before equilibration (green dashed line) and after equilibration (red line), and compared with the FEP computed for equilibrium sampling (blue dotted line).

### Non-equilibrium sampling with multiple short trajectories of fixed length and trap state B

A total of 25000 short trajectories were simulated with the following parameters: diffusion coefficient $D = 1$, maximum number of steps per trajectory $nsteps = 100$, and time step $\Delta t_0 = 0.0005$. Trajectories terminate upon reaching boundary $B$. The short trajectories were concatenated into a single long trajectory, comprising approximately 2000000 steps. Supplementary Fig. S3 shows that the approach accurately computes the committor time-series. The non-equilibrium FEP (green dashed line in Supplementary Fig. S3e) is high around boundary state B due to trajectories terminating upon reaching state B. The equilibrium FEP, obtained by re-weighting the non-equilibrium trajectories, is in agreement with the equilibrium profile shown by the blue dotted line in Supplementary Fig. S3e.

### Comparison of metrics

The figures present precision-recall curves, Kaplan-Meier curves, and time-dependent AUCs. Precision-recall curves are recommended for imbalanced datasets, while Kaplan-Meier curves and time-dependent AUCs are commonly used in survival analysis. Supplementary Fig. S4 shows results for an "ideal system", specifically chosen to highlight cases where all three metrics perform well and distinguish between optimal and sub-optimal RCs, in agreement with other metrics shown in Fig. 4. The precision-recall curve for the optimal RC (Supplementary Fig. S4d) has a higher AUC than that for the sub-optimal RC (Supplementary Fig. S4a). Kaplan-Meier curves for the optimal RC decay more slowly, and the time-dependent AUC is higher than for the sub-optimal RC. For a "typical system" (Supplementary Fig. S5) and a system with imbalanced data (Supplementary

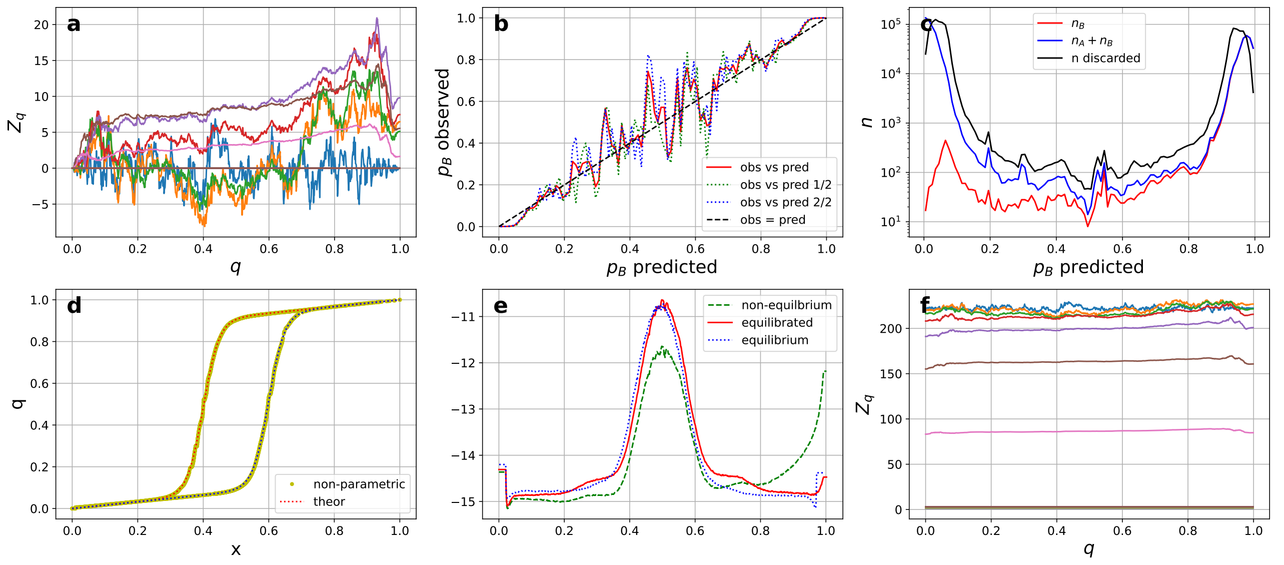


Supplementary Figure S3: Non-equilibrium sampling with multiple short trajectories of fixed length and trap state B. Notations are the same as in Supplementary Fig. S2.

Fig. S6), these metrics perform weakly in differentiating between optimal and sub-optimal RCs for describing dynamics, similar to Figs. 5-6

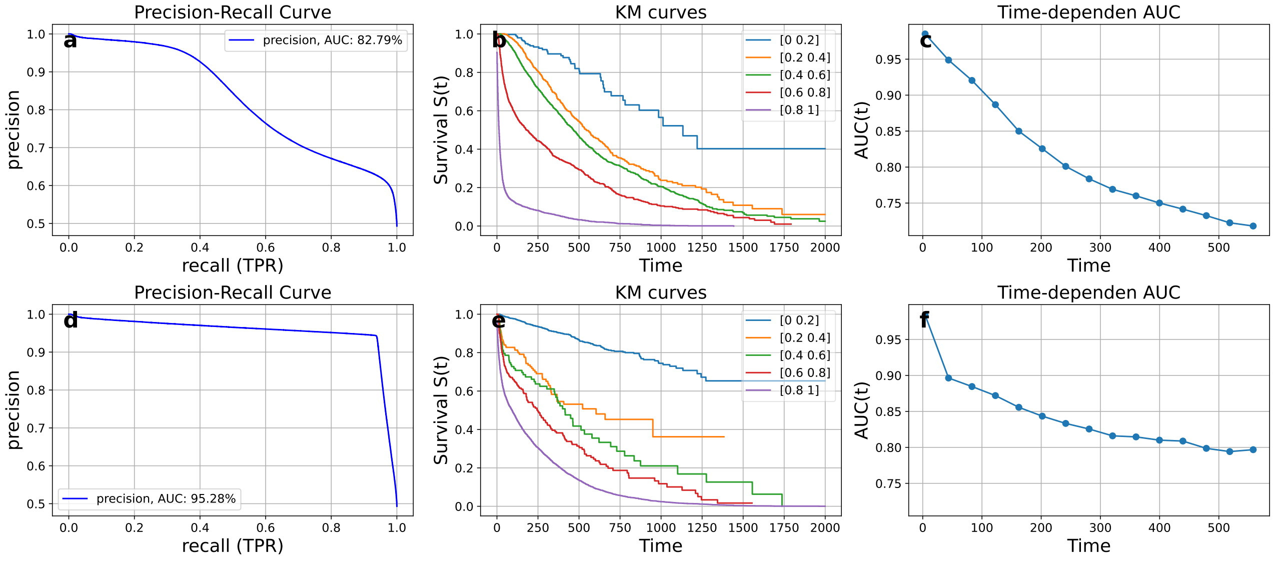


Supplementary Figure S4: Comparison of metrics for the "ideal" system. The first row (panels **a**, **b**, **c**) presents the results for the sub-optimal RC, $q_{sub}$, while the second row (panels **d**, **e**, **f**) shows the results for the optimal committor, $q$. **a**, **d**) Precision-recall curves; **b**, **e**) Kaplan-Meier curves; **c**, **f**) and time-dependent AUCs.

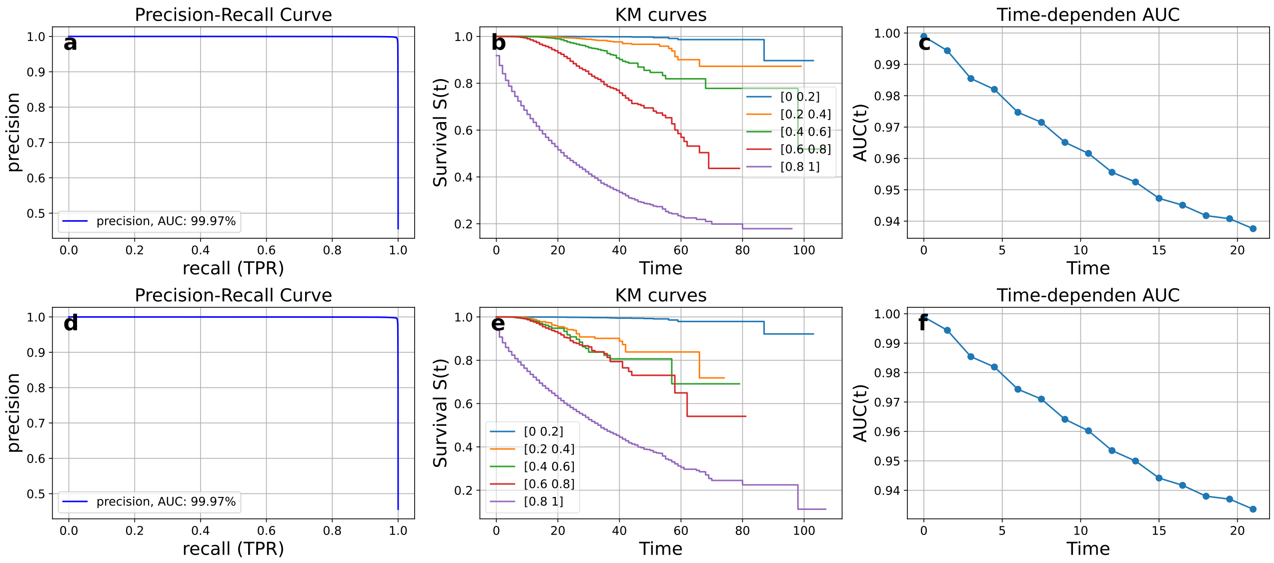


Supplementary Figure S5: Comparison of metrics for the ”typical” system. Notations are the same as in Fig. S4.

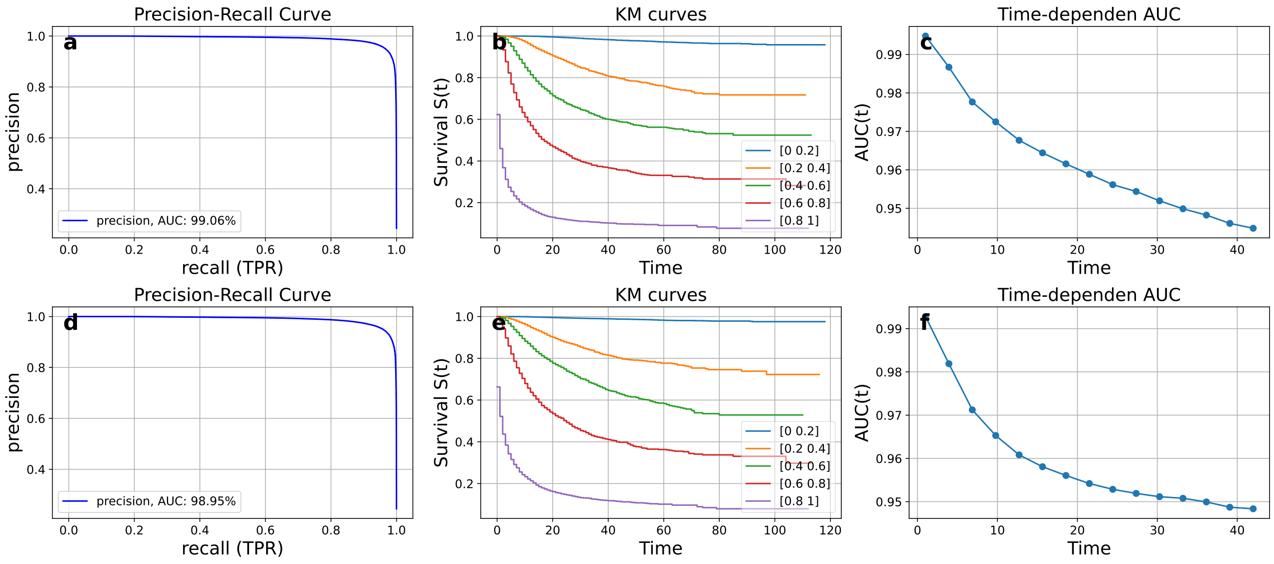


Supplementary Figure S6: Comparison of metrics for the ”imbalanced data” system. Notations are the same as in Fig. S4.
